# Three genomes of Osteoglossidae shed light on ancient teleost evolution

**DOI:** 10.1101/2020.01.19.911958

**Authors:** Shijie Hao, Kai Han, Lingfeng Meng, Xiaoyun Huang, Chengcheng Shi, Mengqi Zhang, Yilin Wang, Qun Liu, Yaolei Zhang, Inge Seim, Xun Xu, Xin Liu, Guangyi Fan

**Affiliations:** School of Future Technology, University of Chinese Academy of Sciences, Beijing, China; BGI-QingDao, BGI-Shenzhen, Qingdao, 266555, China; BGI-Shenzhen, Shenzhen 518083, China; State Key Laboratory of Agricultural Genomics, BGI-Shenzhen, Shenzhen 518083, China; Department of Biotechnology and Biomedicine, Technical University of Denmark, Lyngby, 2800, Denmark; Comparative and Endocrine Biology Laboratory, Translational Research Institute-Institute of Health and Biomedical Innovation, School of Biomedical Sciences, Queensland University of Technology, Brisbane 4102, Queensland, Australia

## Abstract

Osteoglossiformes is a basal clade of teleost, originated from late Jurassic and had seen the process of continental drift. The genomic differences amongst Osteoglossiformes species should reflect the unique evolve history of that time. Here, we presented the chromosome-level genome of *Heterotis niloticus* which is the only omnivore species of Osteoglossidae spreading in Africa. Together with other two Osteoglossidae species genomes of *Arapaima gigas* and *Scleropages formosus* which spread in South America and Australia respectively, we found great evolutionary differences in gene families and transposable elements. Phylogenetic analysis showed that the ancestor of *H. niloticus* and *A. gigas* diverged with *S. formosus* at ∼106.1Mya, consistent with the time of Afro-South American drift and *A. gigas* speciated from the ancestor of *H. niloticus* and *A. gigas* at ∼59.2 Mya, consistent with the separation of Eurasia and North American continents. And we proposed the evolutionary traces of Osteoglossidae species based on comparative genomics analysis and their living geographic habitats. We identified loss of LINEs and LTRs, fast evolutionary rate in parallel to fast pseudogenization rate in *H. niloticus* and *A. gigas* comparing to *S. formosus* during the evolutionary process. We also found notable OR genes contraction in *H. niloticus*, which might be related to the diet transition. Taken together, we reconstructed the evolutionary process of Osteoglossidae using three representative genomes, providing a possible clue for biogeographic and evolution study of ancient teleost clade.

## Introduction

The Osteoglossiformes is a basal clade of teleost which comprise five living groups (Hiodotidae, Osteoglossidae, Pantodontidae, Notopteridae and Mormyroidea). The Osteoglossidae contained two extant clades including Osteoglossinae and Heterotidinae spreading in Asia, America, Africa and Australia[1]. The Osteoglossiformes fossils had been found earliest in the late Jurassic[2]. Therefore, it had witnessed the break-up of Gondwana supercontinent[3-5]. The evolutionary history of Osteoglossiformes species can represent a typical example of biogeography and have been studied extensively through morphological and molecular biological tools[2, 4, 6]. As the rise of genomics, several Osteoglossiformes species had their genomes decoded[7-9]. These works contributed a great amount of genetic information and facilitated the further evolutionary analysis.

Osteoglossidae contains four genera Arapaima, Heterotis, Osteoglossum and Scleropages. The *Heterotis niloticus*, the only omnivore in Osteoglossiformes[10, 11], together with its sister species *Arapaima gigas* and *Scleropages formosus* form a good example to investigate the genetic basis of this ancient teleost clade[12]. *H. niloticus* lives in Africa while *A. gigas* mainly distributes in South America and *S. formosus* distributes in Southeast Asia, characterized by several morphological differences. For example, *A. gigas* is one of the biggest freshwater fish with body size reaching to 2.75 meters while *H. niloticus* and *S. formosus* is only one meter long [13-15]. However, they also show several similar behaviors, such as they all guard larva for a long time[16-18].

Besides, the genome sequences of *S. formosus* and *A. gigas* have been available. In this study, we assembled a chromosome-level *H. niloticus* genome and investigated the genomic differences amongst *A. gigas, H. niloticus* and *S. formosus* genomes, attempting to explore the evolutionary process of Osteoglossidae.

## Results

### *H. niloticus* genome assembly and annotation

The *H. niloticus* genome was sequenced about 144.36 Gb (∼186.42-fold of whole genome) using single tube long fragment reads (stLFR) technology[19] on the BGISEQ-500 sequencing platform (**Supplementary table 1**). We assembled the genome into 4,244 scaffolds which span ∼669.73 Mb (99.45% of the estimated genome size 673.41Mb) with an ultra-long scaffold N50 of ∼9.62 Mb. To improve the continuity of this assembly, we sequenced more 10.17 Gb (∼13.14-fold) single molecular long reads using Nanopore sequencing platform, resulting in a notable increasing of contig N50 value from 255.61 Kb to 2.31 Mb using TGS-GapCloser[20] (**Supplemental table 2**). Based on the high quality draft genome assembly, we also sequenced 21.23 Gb data of a Hi-C library and anchored 647.59 Mb (∼96.80% of the whole assembly) scaffold sequences onto 20 chromosomes (**Fig. 1a, Supplemental table 3**), consistent with the reported *H. niloticus*’s karyotype[11]. To double check the taxonomy information of this sequencing sample, we assembled a complete *H. niloticus* mitochondria genome of 16.55 Kb using MitoZ[21], which was phylogenetically closest to published *H. niloticus* mitochondria genome. We then annotated 24,142 gene models in the genome (**Supplemental table 3**), ∼89.54% of which had homologs in public databases. And ∼97.6% and ∼96.8% of 2,586 vertebrate Benchmarking Universal Single-Copy Orthologs (BUSCO)[22] were completely covered by our genome assembly and gene set, respectively, revealing the high quality of our assembly and annotation.

**Figure 1.**
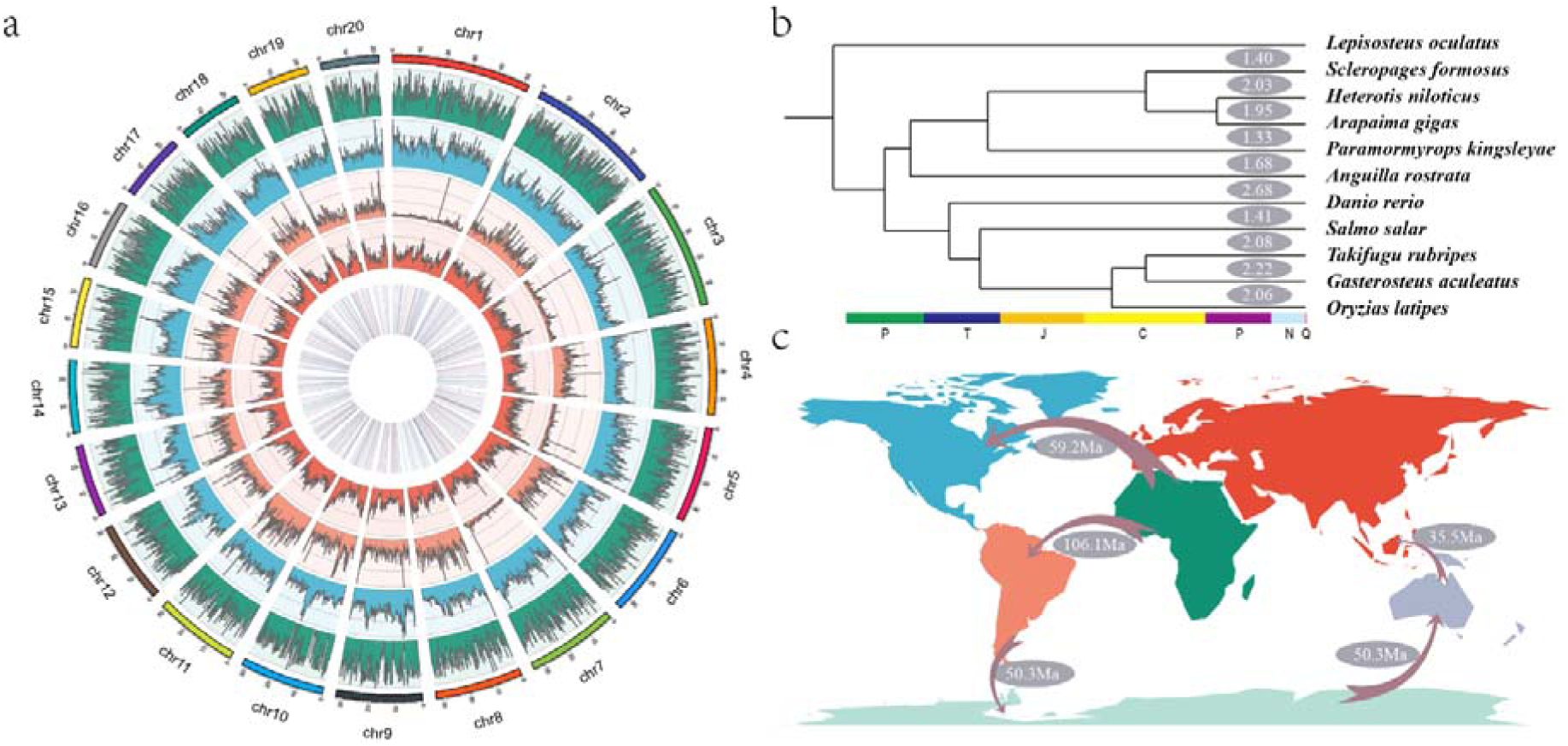
The evolution of *H. niloticus* genome. **a**) Characteristics of the assembled *H. niloticus* genome. The tracks from outer to inner represent gene density, TE density, tandem repeat density, GC content and non-coding RNA respectively. **b**) Phylogenetic relations of 10 teleost fishes with *L. oculatus* as outgroup. The numbers upside clades represent **c**) *A. gigas* and *S. formosus*’ migration pattern through the continents drift.

### Species evolution traces along with the geographic drift

The Osteoglossiformes was speculated that originated from Gondwana supercontinent and spread to modern continents along the continental drift[4]. To investigate the evolutionary process of the Osteoglossidae, we used 355 one-to-one orthologs across ten teleost species, and *Lepisosteus oculatus* as outgroup, to build a phylogenetic tree including *H. niloticus, A. gigas* and *S. formosus* (**Fig. 1b**). Firstly, we calculated the divergence time of *S. formosus* and the common ancestor of *H. niloticus* and *A. gigas* at ∼106.1 Mya (± 20 Mya) [3, 8, 23], which is coincided with the separated time of South America and Africa continents by Afro-South American drift at ∼110 Mya[5]. At the original of separated time, the ancestor of *S. formosus* might live in South America. However, the modern *S. formosus* only live in Southeast Asia[24], suggesting their ancestor might migrated to the current habitats from South America by tectonic-mediated Gondwanan fragmentation (such as the fragmentation of South America–Antarctica–Australia and/or the fragmentation of Southeast Asia–Australia) (**Fig. 1c**).

Besides, we also estimated that *H. niloticus* and *A. gigas* diverged from the common ancestor at ∼59.2 Mya (**Fig. 1b**). The modern *H. niloticus* only lived in African freshwater and the modern *A. gigas* only live in South American freshwater. The land connection between North America and Eurasia still existed ∼60Mya at least[25]. Thus, these results suggest the common ancestor of *H. niloticus* and *A. gigas* might migrated to South America through the path of Africa-Eurasia-North America-South America before ∼60Mya. Then *H. niloticus* and *A.gigas* speciated because of the separation of North America and Eurasia (**Fig. 1c**).

### Fast evolution of *H. niloticus* and *A. gigas* lineage

The total length of this genome assembly (∼669 Mb) is very close to that of *A. gigas* (∼665 Mb), but is notable less than that of *S. formosus* (∼779 Mb). To trace the process of the genome size change between *S. formosus* and the other two genomes, we firstly estimated the molecular evolution rates of these three clades by calculating the substitutions accumulate values in 1849 one-to-one ortholog genes of *A. gigas, H. niloticus, S. formosus* and *L. oculatus*[26]. We found the mean dN and dS (The number of non-synonymous and synonymous substitutions per site) values of *S. formosus* (0.12 and 1.28) are both lower than those of *H. niloticus* (0.15 and 1.88) and *A. gigas* (0.15 and 1.80), suggesting HA lineage (*H. niloticus* and *A. gigas*) had a faster lineage-specific variation rate than *S. formosus* (**Fig. 1b**). We also identified the syntenic blocks in intra-*H. niloticus* genome, intra-*S. formosus* genome and inter-*H. niloticus-S. formosus* genomes[27]. We found there are distinct collinear relationships of inter-*H. niloticus-S. formosus*, illustrating they had experienced slight chromosome structure variation after their speciation (**Fig. 2a**). Interestingly, we are able to see a more unambiguous syntenic blocks in intra-*S. formosus* than intra-*H. niloticus* chromosomes, also indicating *H. niloticus* genome has a faster evolutionary rate resulting in the relatively chaos pair-wise relationship of their common TS-WGD events (∼350 Mya) in *H. niloticus* genome[28] (**Fig. 2b&2c**).

**Figure 2.**
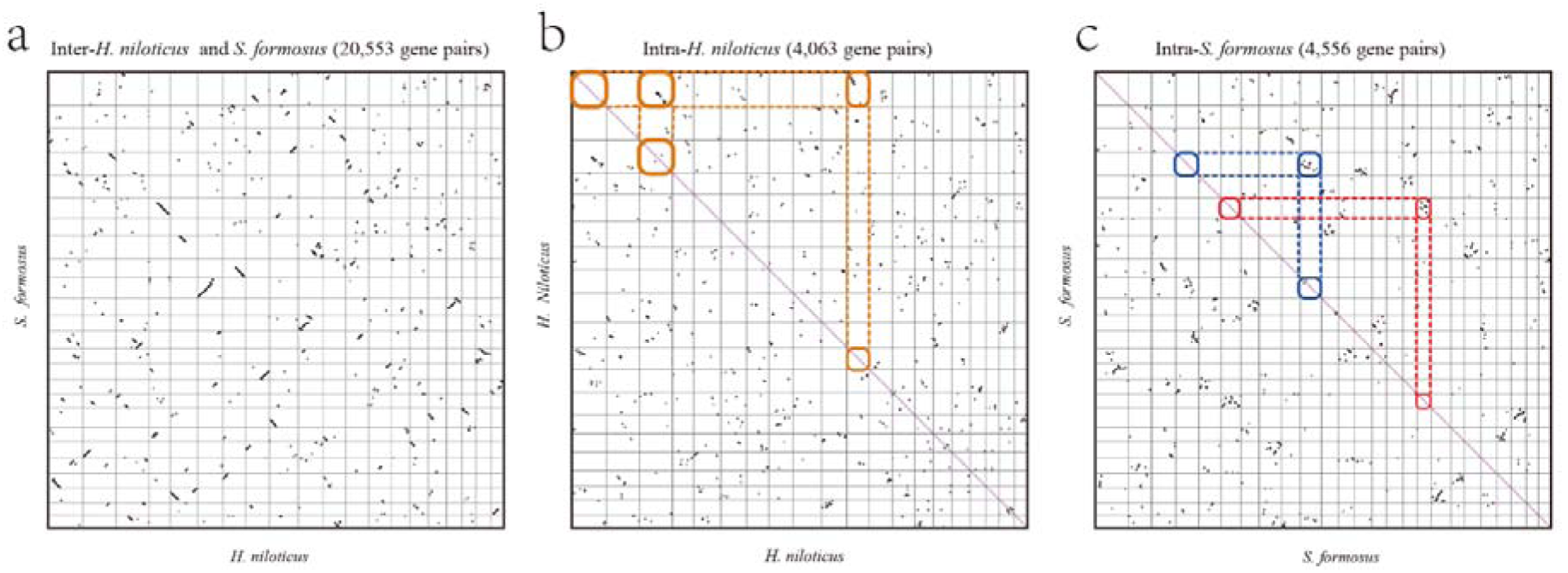
Synteny patterns of *H. niloticus* and *S. formosus*. **a**) Synteny pattern of *H. niloticus* and *S. formosus*. **b**) Synteny pattern of intra-*H. niloticus*. **c**) Synteny pattern of intra-*S. formosus.*

### The loss of TE in HA lineage

Transposable elements (TE) play a key role in genome evolution. It’s reported that TE can regulate gene expression[29], serve as raw material for new gene[30], reshuffle coding sequences (CDS)[31] and generally proportional to genome size[32]. The proportion of TE sequences in *H. niloticus* (18.74%) and *A. gigas* (18.16%) are similar, but notable less than that of *S. formosus* (29.51%) (**Supplemental table 4**). Among them, two TE subtypes of LINE and LTR are notable different between *S. formosus* (LINE: 16.36% and LTR: 12.33%) and HA lineage (LINE: ∼4.4% and LTR: ∼3.84%) (**Fig. 3a**). In detail, we extracted common classes of TEs and built phylogenetic trees to investigate their evolutionary relations. Many of the TEs in three species were classified into closest clades, such as DNA/TcMar, DNA/Crypton-V and DNA/hAT-Tol2 (**Fig. 3b-3d**). Interestingly, LINE/LINE and LTR in *S. formosus* are significantly more abundant than in *H. niloticus* and *A. gigas* (**Fig. 3e** & **3f**). Because *S. formosus* had a relative slow evolutionary rate after experiencing WGD event, we proposed *H. niloticus* and *A. gigas* simplify the genome size by loss of LINEs and LTRs from their genome after diverging with *S. formosus*, rather than *S. formosus* experience species-specific TE expansion.

**Figure 3.**
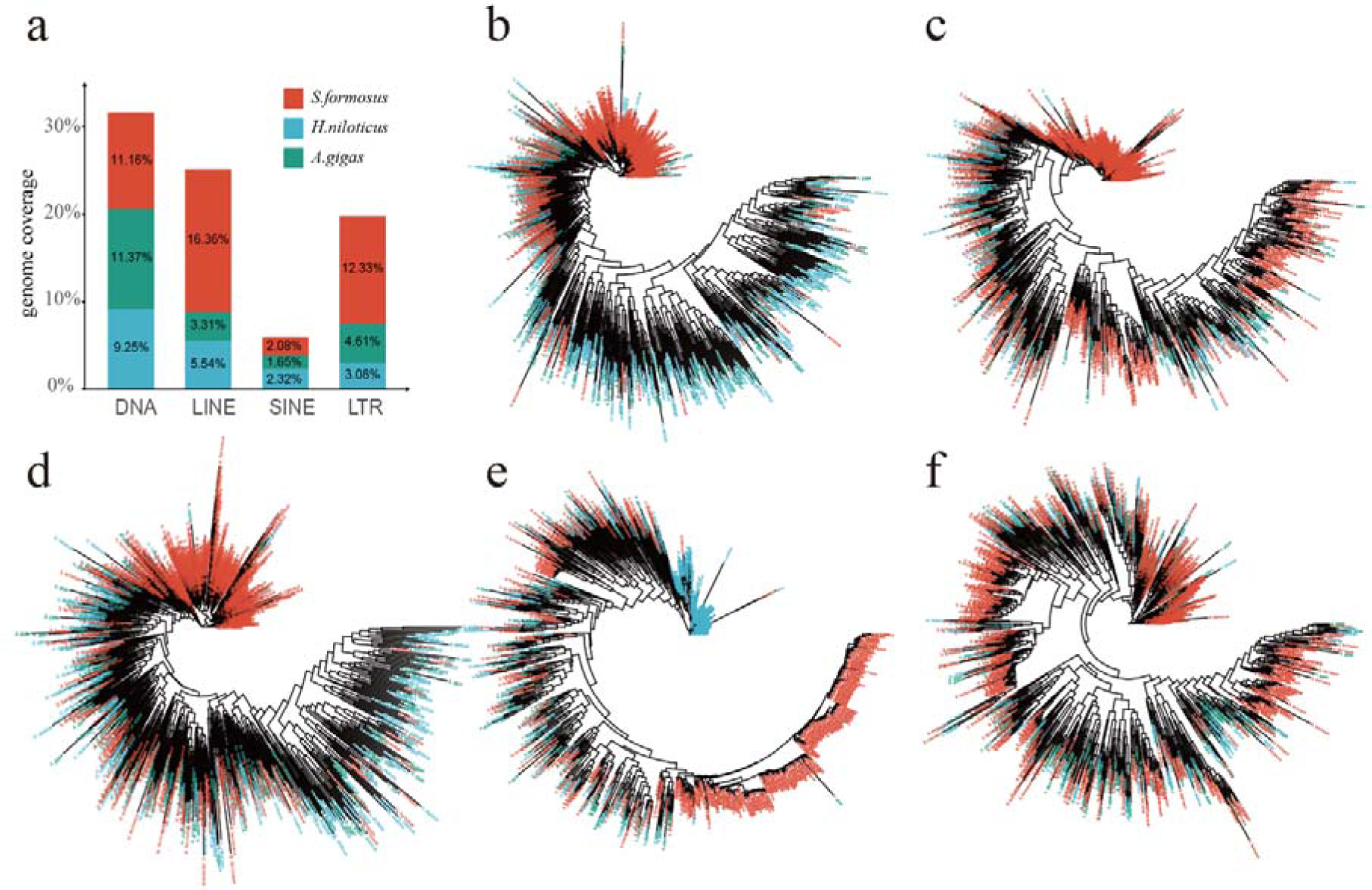
*A. gigas, H. niloticus* and *S. formosus*’ transposable elements content and phylogenetic relations of same TE class. **a**) TE contents of 3 bonytongue fish genomes. **b-f**) Phylogenetic relations of DNA/TcMar, DNA/Crypton-V, DNA/hAT-Tol2, LINE/LINE and LTR.

### The higher pseudogenization rate of HA lineage

Gene duplications are always related with the physiologic functions of organisms, such as environment adaptation[33]. However, during the evolution process, duplicated genes faced less selective stress, and some of them were malfunctioned rapidly by pseudogenization, became to the “genomics fossils”[34]. Therefore, we identified the pseudogenes in these three genomes. Expectedly, we detected 17,311 and 22,712 pseudogenes in *H. niloticus* and *A. gigas* respectively, which is notable more than 9,254 pseudogenes in *S. formosus*, suggesting *H. niloticus* and *A. gigas* have higher pseudogenization rate than *S. formosus*, consistent with the evolutionary rate. There were 4,653 pseudogene families shared by *H. niloticus* and *A. gigas*, but all members of these families are retained functional in *S. formosus* (**Fig. 4a**). The functional enrichment analysis of these pseudogenes showed they significantly enriched in metabolic pathways, synthesis and degradation of ketone bodies, endocytosis and etc. pathways. Interestingly, we found 1,127 *H. niloticus* specific pseudogene families functional enriched in salivary secretion, olfactory transduction and etc. which might be correlated with its diet behaviors (**Fig. 4b**).

**Figure 4.**
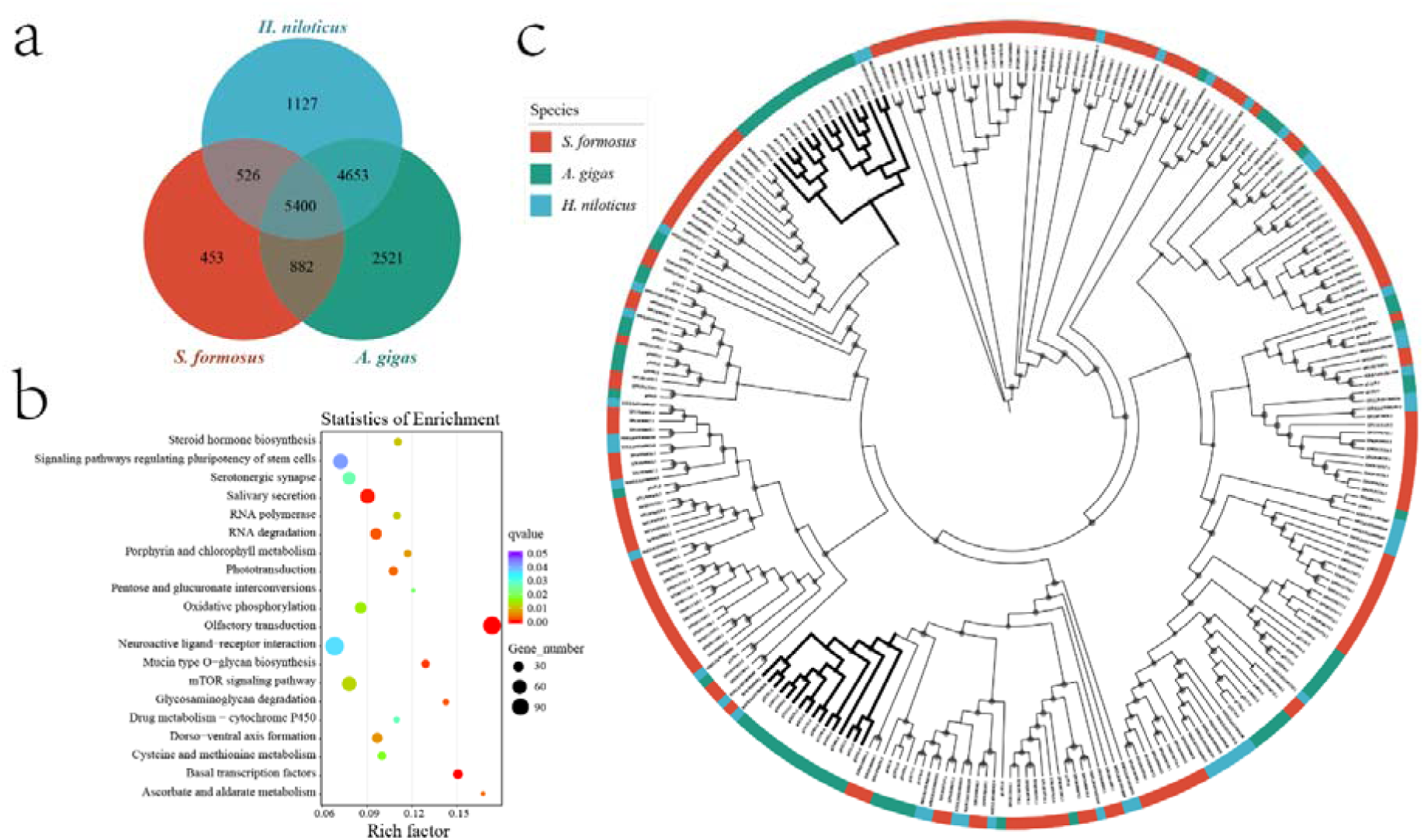
Comparisons of the pseudogenes among three genomes. **a**) Three bonytongue fishes’ pseudogene related gene family’s distribution. **b**) The *H. niloticus* specific pseudogenes KEGG enrichment results. **c**) Gene tree of three bonytongue fishes’ olfactory receptors.

Pseudogenes genesis is correlated with the birth of gene families[35]. And expanded or contracted gene families always be related with physiologic function as reported[36, 37]. We detected 1,210, 424 and 829 expanded gene families in *A. gigas, H. niloticus* and *S. formosus* genome respectively. The KEGG functional enrichment results showed that *A. gigas* and *S. formosus* experienced significantly expansion in 14 pathways such as Olfactory transduction, Salivary secretion, Cell adhesion molecules (CAMs), NOD-like receptor signaling pathway and etc. which were not found in *H. niloticus* (**Supplemental table 5**, *p*-value<0.01 and *q*-value<0.01). And these three species only share three significant expanded pathways, including Gap junction, Necroptosis and NOD-like receptor signaling pathway.

### Genes related to diet transition analysis

*H. niloticus* is a kind of omnivore which have a wide range of preys such as small benthic fishes, shrimps, plants and insects[10, 16] while its sister species, *S. formosus* and *A. gigas*, are predominantly dependent on fish preys[14, 38]. We comprehensively examined the taste receptors of all tastes including sweet, umami, bitter, sour and salty in these three genomes and found no significantly biased expansion or contraction in them (**Supplemental table 6**). The vertebrates have three kinds of odorant receptors including olfactory receptors (OR), vomeronasal receptors V1R and V2R[39]. We also implemented odorant receptors comparative analysis of these three genomes. Interestingly, we found OR genes (K04257) were significantly contracted in *H. niloticus* (40) comparing to *A. gigas* (70) and *S. formosus* (160). Through the gene tree of olfactory receptors (**Fig. 4c**), we observed that *S. formosus* kept relative complete gene copies in most clades and *A. gigas* experienced contraction in several clades except for two clades (marked by bold clades). Besides, there were 14, 15, 14 V1R genes and 2, 2, 1 V2R genes in *A. gigas, H. niloticus* and *S. formosus* respectively which showed no notable phylogenetic expansion (**Supplemental table 7**). Therefore, we speculated that the OR gene contraction may play a key role in diet transition of *H. niloticus*.

## Discussion

The *H. niloticus, A. gigas* and *S. formosus* located at Osteoglossidae’s two clade, which spread in South America, Africa, Asia because of continental drift and had experienced different environmental changes. Therefore, the genomic differences amongst them should implicate the evolutionary processes. Here, we presented *H. niloticus* genome and implemented comparative genomic analysis amongst these three species and found the loss of LINEs and LTRs, high pseudogenization rate and poor syntenic relation in *H. niloticus* and *A. gigas* comparing to *S. formosus*. All these results indicated a faster evolutionary rate of HA lineage than *S. formosus*. The phylogenetic history of Osteoglossida is an interesting topic and we firstly constructed the phylogenetic relation of three Osteoglossida species using genomic tools, completing the biogeographic pattern of Osteoglossida. Moreover, the functional enrichment results of families that experienced pseudogenization of *H. niloticus* present a great coincidence with the gene families expansion analysis of other two species in olfactory transduction and salivary secretion pathways. Therefore, OR gene contraction and pseudogenization of olfactory transduction and salivary secretion pathways may be the reasons of diet transition.

However, the data we provided is not sufficient for all problems we put forward. More researches are required in future such as the fossils evidence searching in Africa and South America. The impact of TEs loss on new gene genesis and gene expression regulation in *H. niloticus* and *A. gigas* also need further deep study. The diet transition of *H. niloticus* and the inter-continental emigration of bonytongues should be transferable environments of fresh water fishes particularly for the fishes live in the same period and similar environment with Osteoglossidae. Moreover, the additional *de novo* genomic researches in Osteoglossidae and other fish will help us to understand the evolution of this ancient teleost clade, such as 10000 fish genome (Fish10K) project.

## Methods

### Sample collection, library construction and genome sequencing

Individual African arowana fish from a seafood market at Xiamen, Fujian province, southeast China was collected and the muscle tissues was used for DNA extraction using the conventional salting-out method. The high molecular weight genomic DNA with average length of 50 Kb was further used to construct a single tube Long Fragment Read (stLFR) library using the MGIEasy stLFR Library Prep kit (PN: 1000005622) according the instructions[40]. Hi-C library was constructed following the Wang’s methods[40] with whole blood tissue of the same individual. Sequencing was conducted on a BGISEQ-500 platform with pair end 100 bp read length. Besides, one nanopore library was also prepared according the instructed protocol using the Oxford Nanopore SQK-LSK109 kit and loaded on the sequencing platform to sequencing.

### Genome survey

The *k*-mer frequencies within the clean stLFR reads were analyzed to estimate major genome characteristics. The occurrences of all 17-mers within both strands were counted using jellyfish v2.2.7[41], and the genome size, heterozygosity as well as repeat content were calculated using GenomeScope[42]. The modeling distribution of 17-mer frequency demonstrated a peak at around 52, with 41,344,762,649 total number of *k*-mers. The estimated haploid genome size was 673.41Mb, of which 31% was inferred to be repeat, low heterozygosity rate (0.13%) was detected though no apparent peak indicated heterozygosity in this genome was detected.

### *De novo* genome assembly

Draft genome sequence was first assembled using Supernova v 2.1.1[43] software and processed with one round of gap closing using Gapcloser v1.12[44] with stLFR data. In this process, the stLFR reads were first pre-processed to be compatibly handled by supernova assembler, using the stLFR2Supernova pipeline (https://github.com/BGI-Qingdao/stlfr2supernova_pipeline). Then, we enhanced the draft assembly using TGS-GapCloser pipeline[20] based on the single molecular long reads.

Hi-C data were used to improve the connection integrity of the scaffolds. We first detected all valid pairs of reads using Hic-Pro v2.8.0[45] by mapping clean Hi-C reads to draft genome sequences, and the valid read pairs were extracted and aligned to the genome using Juicer v1.5[46]. Then the assembled fragments of DNA were ordered and oriented using 3D-DNA pipeline[47] based on the Juicer Hi-C contacts (‘merged_nodups.txt’ file). Manual review and refinement were also performed by using Juicebox Assembly Tools v1.9.0[48] to identify and remove the remaining assembly errors.

### Genome annotation

We detected and annotated repetitive sequences, mainly tandem repeats (TRFs) and transposable elements (TEs), in the genomes. For the annotation of TRFs, Tandem Repeats Finder v 4.04 program[49] was employed. The TEs were annotated by a combination of both *de novo* prediction and homology-based identification. Briefly, the genome sequences were first *de novo* searched using LTR_Finder[50] and RepeatModeler[51] to find sequence elements with specific consensus models of putative interspersed repeats. The non-redundant self-contained repeat library was then searched against the genome using RepeatMasker[51]. In the homology-based detection, the genome sequences were aligned to both the public Repbase 21.01 and transposable element protein database (included in the RepeatMasker package) to detect interspersed repeats. Evidences including ab initio gene predictors and homology to proteins previously discovered in other sequenced genomes as well as transcript sequences were integrated together to make a comprehensive gene structure prediction. Augustus[52], GlimmerHMM[53] and Genescan[54] were applied for ab initio gene finding with best parameters trained for zebrafish and/or vertebrates. For homology-based prediction, nonredundant protein sequences from 5 species (*Oreochromis niloticus, Pundamilia nyererei, Maylandia zebra, Astatotilapia calliptera* and *Scleropages formosus*) were aligned against African arowana genome using GeneWise v2.4.1 program[55]. Furthermore, transcript sequences were constructed based on the RNA-Seq alignment to the genome that generated by using HISAT, and candidate coding regions within the transcripts were further detected, in which ORFs with homology to known proteins were also identified via blast (against SwissProt database) and pfam searches, using TransDecoder v5.5.0 (https://transdecoder.github.io/). Final consensus gene models were produced by integrating those disparate sources of gene structure evidence using GLEAN software[56]. 24,146 genes, covering 96.8% vertebrate BUSCOs, were predicted in the African arowana genome with average length 14911.23 bp. The length distributions of mRNA, coding sequences, exon and intron were close similar to that of related species.

Functional annotations of the predicted genes were performed by aligning protein sequences to KEGG release 84.0, Swissprot release 201709, Trembl release 201709 and Clusters of Orthologous Groups (COGs) database. The results show that 21,609 (89.49%) protein-coding genes were assigned successfully to at least one well-modelled functional category.

### Evolutionary phylogeny of African arowana

To reveal the phylogenetic relationships of African arowana and other bony fishes, gene set of five Clupeocephala species (*Danio rerio, Salmo salar, Oryzias latipes, Gasterosteus aculeatus* and *Takifugu rubripes*), one Elopomorpha species (*Anguilla rostrate*) and three Osteoglossomorpha species (*Scleropages formosus, Paramormyrops kingsleyae* and *Arapaima gigas*), plus one species from Lepisosteiformes (*Lepisosteus oculatus*) as outgroup, were downloaded from NCBI and further used to detect gene clusters. We extract the longest transcript from unique genomic loci to eliminate redundant splicing, and retained coding sequences longer than 150 bp from each dataset to discard possibly unreliable gene predictions. We performed all-versus-all BLAST search for protein sequences of these 11 species and the resultant matches were sorted out for filtering redundant and segments, then the genes were further clustered using hcluster_sg tool (https://sourceforge.net/p/treesoft/code/HEAD/tree/branches/lh3/). The genes were grouped into 23654 clusters, of which 355 were single-copy.

We performed multiple sequences alignment using MUSCLE v3.7 software[57] for each gene and further concatenated the alignments into super-matrix. Phylogenetic relationships of these species were inferred using MrBayes v3.1.2[58] based on the fourfold degenerate site of the supergene. Divergence time of our target species were also determined using MCMCTree[59] with the prior timelines from TimeTree[60] as calibrations. Given the phylogenetic relationship and divergence time, we analyzed the changes in gene family size using CAFE v2.1[61]. We compared the gene pairs in the paralogous and orthologous families detected by using wgd v1.0.1 package[62], the distribution of synonymous mutation rate (*Ks*) was used as indicator of the duplication and divergence event in three Osteoglossidae species (*Scleropages formosus, Arapaima gigas* and *Heterotis niloticus*).

### Timeline of pseudogene in Osteoglossidae

We scanned matches of the protein sequences against the genomes of Osteoglossidae species (*Scleropages formosus, Arapaima gigas* and *Heterotis niloticus*) and detected possible pseudogenes separately using PseudoPipe[63] annotation tool. Pseudogenes overlapping genes and/or repeats were filtered. In the analysis of homology, pseudogenes were assigned into different gene families based on the clustering of their parent genes. We also aligned the pseudogenes to their parent genes and used the synonymous substitution rate (*Ks*) as indicator of pseudogene age.

### Genes and transposable elements phylogenetic trees construction

We investigated the phylogenetic relations with 3 fishes’ repeats of same class and genes of same function according to the results of RepeatMasker and KEGG v84.0 (https://www.kegg.jp/) annotations respectively. The gene trees with species specific expanded clades, of which nodes number was more than 10 and any species’ genes occupied more than 80%, were extracted with in-house python script and ete3[64] module. All the gene trees and repeat trees were built by FastTree[65] and visualized by ggtree[66] and iTOL[67].

## Author contributions

Shijie Hao and Guangyi Fan conceived and designed the study. Mengqi Zhang and Yilin Wang performed sample collection and sequencing. Xiaoyun Huang performed assembly. Lingfeng Meng performed genome annotation and partial phylogenetic analysis. Kai Han performed pseudogene related analysis and partial phylogenetic analysis and designed the figures. Shijie Hao wrote the manuscript. Guangyi Fan and all other authors revised and read the manuscript.

## Acknowledgements

This work is supported by the special funding of “Blue granary” scientific and technological innovation of China (2018YFD0900301-05). The work also received the technical support from China National Gene Bank.

